# Gene structure, differential exon usage, and expression of the testis long intergenic non-protein coding RNA 1016 in humans reveals isoform-specific roles in controlling biological processes

**DOI:** 10.1101/2021.03.26.437262

**Authors:** Enrique I. Ramos, Barbara Yang, Yasmin M. Vasquez, Ken Y. Lin, Ramesh Choudhari, Shrikanth S. Gadad

## Abstract

Long noncoding RNAs (lncRNAs) have emerged as critical regulators of biological processes. The constant expansion of newly-identified lncRNA genes requires that each one be comprehensively annotated to understand its molecular functions. Here, we describe a detailed characterization of the gene which encodes long intergenic non-protein coding RNA 01016 (*LINC01016*, a.k.a., *LncRNA1195*) with a focus on its structure, exon usage, and expression in human and macaque tissues. In this study, we show that it is exclusively conserved among non-human primates, suggesting its recent evolution and is expressed and processed into 12 distinct RNAs in testis, cervix, and uterus tissues. Further, we integrate *de novo* annotation of expressed *LINC01016* transcripts and isoform-dependent gene expression analyses to show that human *LINC01016* is a multi-exon gene, processed through differential exon usage with isoform-specific functions. Furthermore, in gynecological cancers, such as cervical squamous cell carcinoma and uterine corpus endometrial carcinoma, *LINC01016* is downregulated; however, its higher expression is predictive of relapse-free survival in these cancers. Collectively, these analyses reveal that, unlike coding RNAs, lncRNA isoforms are differentially regulated and precisely processed in specific tissues to perform distinct biological roles.

**One sentence summary:** The distinct molecular role of *LINC01016* isoforms reveals intricate biology associated with lncRNA transcription and processing.

## Introduction

Long noncoding RNAs (lncRNAs) play critical roles in normal physiological processes and aberrant conditions, such as cancer (1,2). Similar to protein-coding messenger RNAs (mRNAs), lncRNAs are transcribed by RNA polymerase II, spliced, polyadenylated, and capped. However, most lncRNAs lack protein-coding potential (1-3), with a few exceptions. LncRNAs are diverse due to different modes of transcription, and they are expressed in a tissue-specific manner. Moreover, lncRNAs have been demonstrated to be excellent tumor biomarkers with therapeutic value (2,4,5); however, their therapeutic application has been limited by the lack of a mechanistic understanding of their actions in the cell. This knowledge gap is due in part to the limited or poor annotation of lncRNAs, which hinders functional studies (6). Thus, it is imperative to determine the gene structure, exon usage, and expression pattern of developmental and disease-relevant lncRNAs to facilitate their functional characterization.

Unlike with protein-coding mRNAs, the annotation of lncRNAs is challenging due to low copy numbers and extreme tissue specificity. Further, it is more difficult to determine the 5’/3’ ends of lncRNAs that are chromatin-associated, transcribed from a divergent promoter, or transcribed in an antisense direction since they coexist with unprocessed transcripts (3,6,7). These challenges have been partly addressed by recent advances in next-generation sequencing technologies, which provide unparalleled prospects in elucidating gene structure, exon usage, expression, isoform-specific biological roles, and evolution of specific noncoding genes (3,8-10). This information can be accessed through publicly available genome and gene expression databases (9,11-13). Notably, a recent study has identified 15,000 to 35,000 differentially expressed lncRNAs across different organs and species, emphasizing the cell-, evolution-, and developmental-specific transcriptional programs (14). However, these studies are insufficient to indicate the tissue-specific isoform functions, and given the nature of lncRNA-dependent-biology, unlike proteins where RNA itself plays a central role. Each lncRNA gene requires isoform-specific annotation and characterization in target tissues to understand its biological role (2).

In this study, we functionally characterize the long intergenic non-protein coding RNA 01016 (*LINC01016*), an intergenic lncRNA, using gene expression data from overexpression of major isoforms, as well as data presented in the Genotype-Tissue Expression (GTEx) project, the Cancer Genome Atlas (TCGA), and the Nonhuman Primate Reference Transcriptome Resource (NHPRTR) (15). We have leveraged these datasets generated through next-generation sequencing to annotate and study the expression, differential exon usage, and isoform-specific roles of *LINC01016* in human and non-human primates. We have interrogated its association with clinical outcome in cervical squamous cell carcinoma, testicular germ cell, and uterine corpus endometrial carcinoma patients.

Results show that the *LINC01016* gene is transcribed using differential exon usage and alternative splicing and, also, different isoforms control specific biological processes. *LINC01016* is abundantly expressed in the testis in the late stages of tissue development, and its gene structure is conserved among non-human primate species. Intriguingly, expression and survival analysis in cervical squamous cell carcinoma, testicular germ cell, and uterine corpus endometrial carcinoma cancer patients suggests that *LINC01016* isoforms are differentially expressed and that higher levels of the aggregate expression of *LINC01016* predict outcome in cervical and testicular cancer patients. Overall, these findings have revealed an essential aspect of lncRNA biology that is rarely associated with protein-coding RNAs.

## Materials and methods

### Database searches and analyses

Primate genomic databases were accessed from the Ensembl Genome Browser. Searches were performed as described elsewhere (16,17). Briefly, BlastN was employed under optimal parameters by means of probes: human *LINC01016* DNA segments (Homo sapiens genome assembly GRCh38.p13). The following genome assemblies were examined: chimpanzee (Pan troglodytes, Pan_tro_3.0), gorilla (Gorilla gorilla, gorGor4), macaque (Macaca mulatta, Mmul_10), marmoset (Callithrix jacchus, ASM275486v1), and mouse lemur (Microcebus murinus, Mmur_3.0).

### RNA-Seq expression datasets for testes in rhesus macaque

NGS paired-end reads (101bp) were downloaded from the NHPRTR (15). The datasets are of Indian-origin rhesus macaque (University of Washington) from testes samples. FASTQ files were downloaded and analyzed by a standard RNA-Seq analysis pipeline.

### *LINC01016* Assembly of Transcriptome Data and Differential Gene Expression

Paired-end reads (150 bp) were mapped to the human transcriptome (hg38) release 101 version (August 2020) by standard RNA-Seq analysis pipeline and differential gene expression done by DESeq2.

### Tumor sample analysis

The expression of *LINC01016* was determined using RNA-seq counts from the GTEx Portal (Version 8) and the TCGA GDC v28. in testicular germ cell tumor (TGCT), cervical squamous cell carcinoma, endocervical adenocarcinoma (CESC), and uterine corpus endometrial carcinoma (UCEC) samples.

### Survival analysis of ovarian cancer, testicular germ cell tumor, and uterine corpus endometrial carcinoma patients

To evaluate the prognostic value of *LINC01016*, we explored its expression in patient samples of cervical squamous cell carcinoma, testicular germ cell, and uterine corpus endometrial carcinoma cancer. The Kaplan-Meier plotter tool was used to plot relapse-free survival for *LINC01016* (19).

## Results

### The human *LINC01016* gene

The *LINC01016* gene is found on chromosome 6 of the human genome, located at the short arm of the chromosome (6p21.31). In human genome assembly, GRCh38.p13, Ensembl genome browser predicts 4 exons of *LINC01016*, spanning 29,408 base pairs between genomic coordinates 33,896,914 and 33,867,506 (Fig. 1A). According to Ensembl (version 101), the *LINC01016* gene is transcribed and processed into 12 different RNA species via a differential arrangement of exons and alternative splicing (Fig. 1B), and each transcript is composed of either one, two, or three exons (Fig. 1B). The sequences for *LINC01016* that are found in the NCBI nucleotide database (accession numbers NR_038989.1 and AK057709) do not contain the entire list of sequences found in Ensembl; however, they are similar to transcript 201 (Fig. 1B).

**Figure 1.**
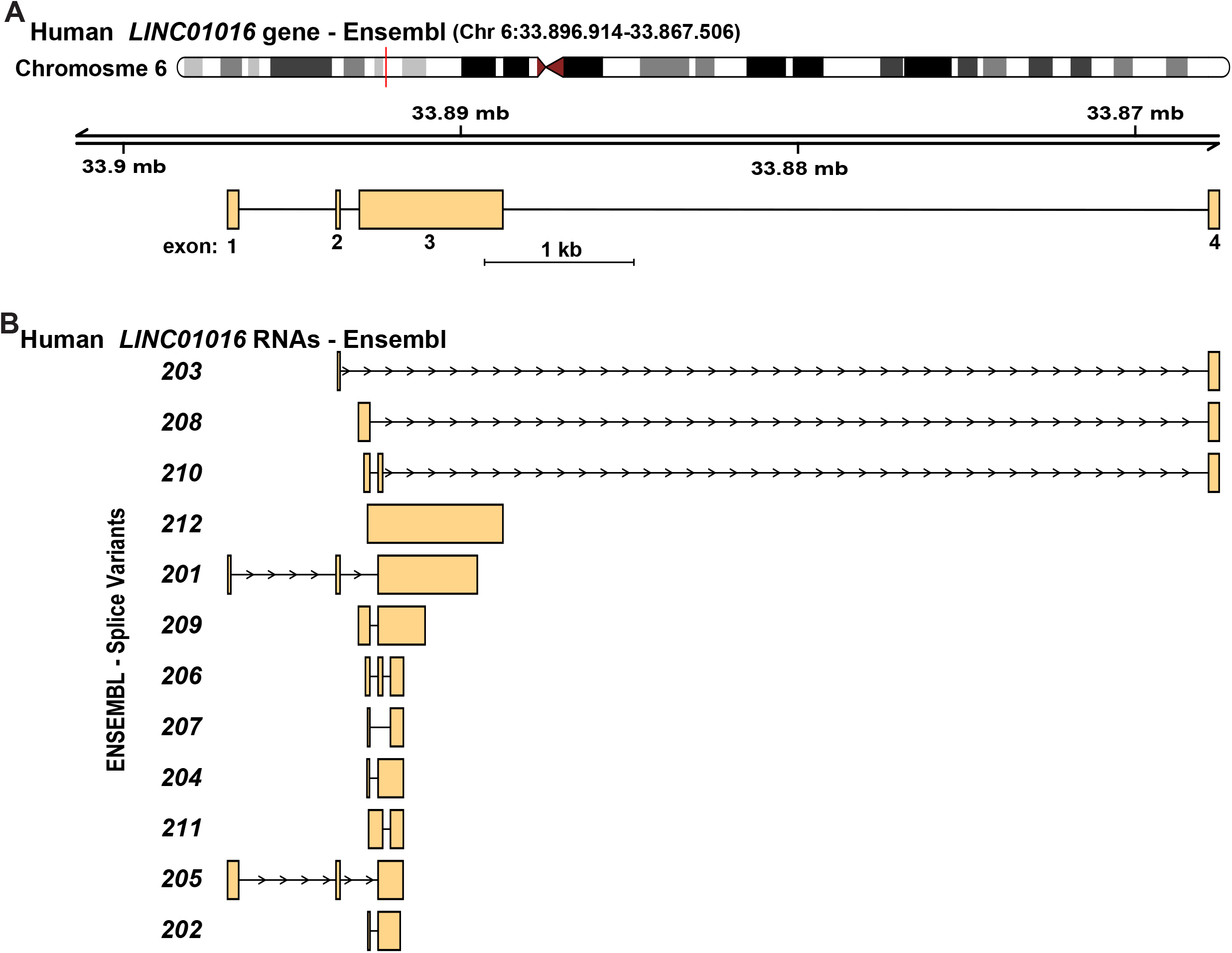
The human *LINC01016* gene in the Ensembl genome database. **A**. Diagram of the human *LINC01016* gene on chromosome 6 from Ensembl. Exons are depicted as boxes and introns as lines. Chromosomal coordinates and a scale bar are shown. **B**. Diagrams of human *LINC01016* RNAs from Ensembl. The exons or portions of exons found in each transcript are illustrated and are aligned with their location within the gene. The scale is the same as in A.

### The *LINC01016* gene in non-human primates

In Ensembl, the *LINC01016* gene has not been annotated in any known non-human primate species, which encouraged us to investigate its presence and location in the target non-human primate genomes. Through examining relevant genomic databases, the *LINC01016* sequence was discovered in 5 different primate species (Fig. 2A and 2B, Table 1), in a region analogous to the location of human *LINC01016*, except for in marmoset, where the chromosome annotation was absent, and in macaque, where it was found on chromosome 4 (Fig. 2B). The data showed the *LINC01016* of interest is comprised of 4 exons (Fig. 2B, Table 1), similarly to human *LINC01016* (Fig. 2A). There was also a similarity in the predicted *LINC01016* gene structure in these primates, and significant DNA conservation was shown for all putative exons, especially in chimpanzee and gorilla (99.37 to 99.19%, respectively) (Fig. 2B, Table 1). However, mouse lemur and marmoset were exceptions, as segments of exon 3 deviated from the other species (Fig. 2B, Table 1). Moreover, RNA-Sequencing analyses showed that *LINC01016* RNAs were present in the testis of macaque species, and an isoform with three exons was predominantly expressed (Fig. 2C). The overall organization of the *LINC01016* gene structure is similar to other reproductive tissue-specific genes, such as *MIR503HG* (7).

**Figure 2.**
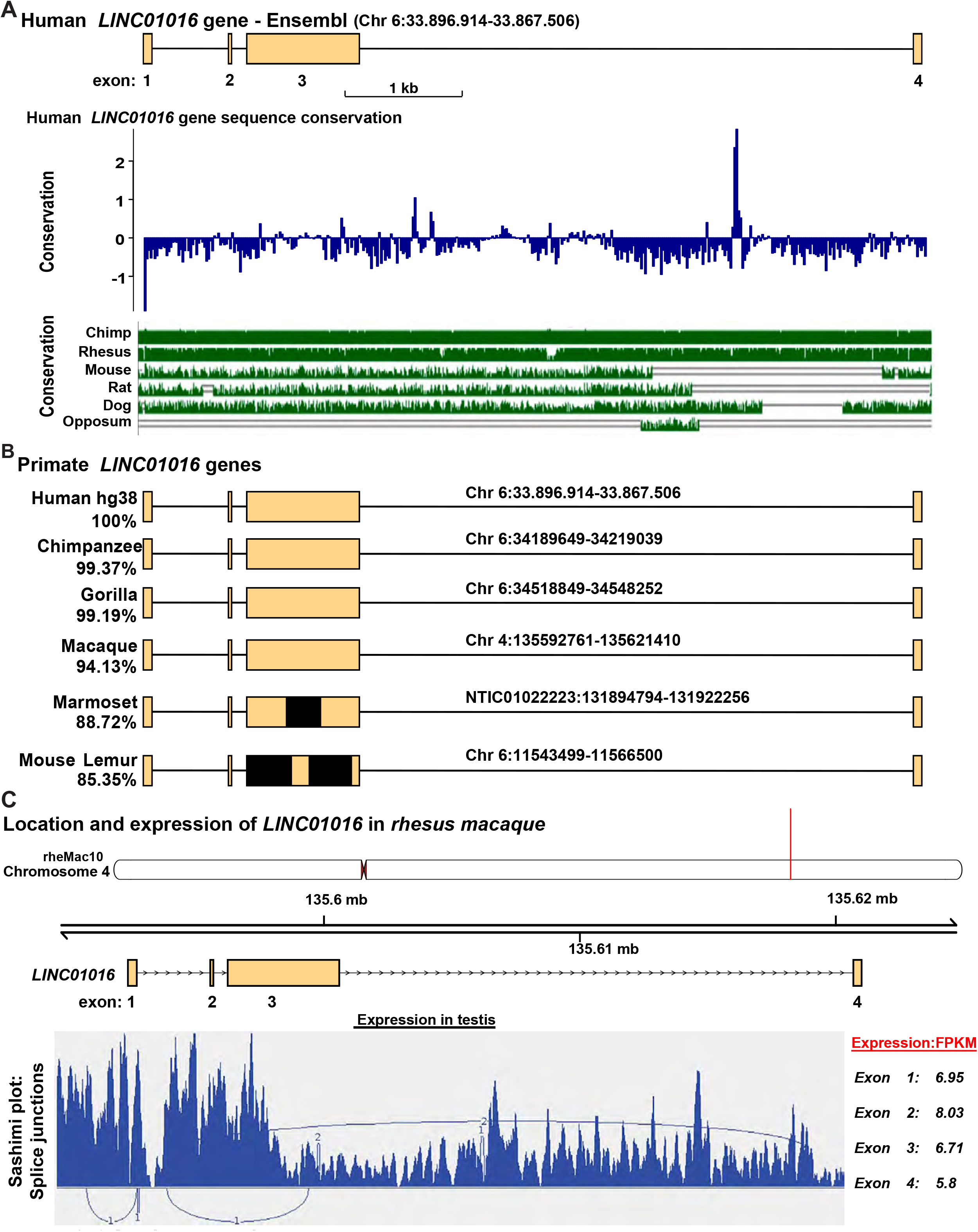
*LINC01016* gene structure and expression in primates. **A**. Human *LINC01016* gene conservation between different vertebrates. Top panel (blue histogram track) shows 100 vertebrate Basewise conservation by PhyloP score. The bottom panel (green alignment track) shows Multiz Alignment & Conservation for 7 vertebrate species. **B**. Conserved identity between different primates and *LINC01016*. Depicted are the predicted genomic structures with conserved exons, and in black are genomic regions missing from BLAST alignments to their respective human sequence. Percentage identity from BLAST search results is provided for each of the species. **C**. The top panel shows the genomic location of the predicted structure for *LINC01016* in rhesus macaque with four conserved exons, similar to human RNAseq experiments. Data from the Nonhuman Primate Reference Transcriptome Resource (NHPRTR) project are shown for expression in testis, and the expression of different exons are shown as FPKM (Fragments Per Kilobase of transcript per Million mapped reads).

**Table 1:**
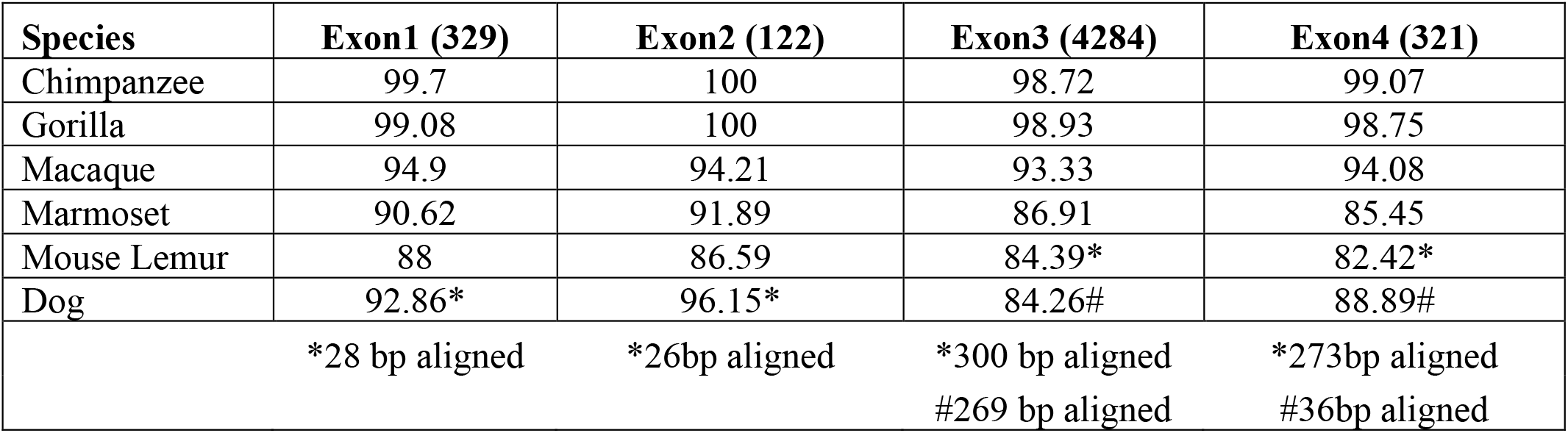
Nucleotide Identity with Human *LINC01016* Exons.

### Expression of *LINC01016* in different human tissues

We sought to understand the extent to which the human *LINC01016* gene had been annotated in Ensembl; analyses were performed by interrogating human gene expression data available from GTEx. Intriguingly, exon-exon junction expression differs across testis, cervix, and uterus tissues (Fig. S1 A and B). In testis, exon-exon junction 5 (Fig. S1 A and B) is predominantly expressed, leading to the formation of a single continuous exon 4 (comprising of exon 3 and 4) (Fig. S1 C), which is similar to isoform 212 described in Ensembl (Fig. S1 B). In contrast, both in the cervix and uterus, junctions 2 and 3 are expressed (Fig. S1 A and B), giving rise to exons 1, 2, and 4 (Fig. S1 C), corresponding precisely to what was defined in the genome database for isoform 201 (Fig. S1 B). However, in the cervix, junction 6 is also expressed but doesn’t affect the exon arrangement (Fig. S1 A and B). Although these data vary from the information presented in Ensembl, they concur firmly with findings recently reported by Sarropoulos *et al*. (14) (see:https://apps.kaessmannlab.org/lncRNA_app/). The observations described above define a human *LINC01016* gene of 2 exons in the testis and 3 exons in both the cervix and uterus (Fig. S1 D).

To understand and determine the differential expression of specific isoforms in respective tumor tissues, we analyzed different *LINC01016* transcripts expression using gene expression profiles from normal (GTEx) and Pan-Cancer (PANCAN) samples. Surprisingly, all the isoforms showed varied expression levels and appeared to be differentially regulated in normal and tumor tissues (cervical, testis, and uterine), suggesting an isoform-specific functional role (Fig S2).

### Human *LINC01016* gene expression

Given the limited evidence of a functional role for the *LINC01016* gene in the literature, the relative abundance of the gene-specific transcripts could indicate its biological significance. Tissue-specific gene expression data were extracted from the GTEx database to determine the abundance of *LINC01016* transcript in healthy human tissues. The analysis revealed that a significant number of *LINC01016* isoforms are expressed in the cervix, testis, and uterus (Fig. S1 D). *LINC01016* transcript levels slightly varied over a 40 TPM range between tissues; the highest expression was seen in testis tissue (Fig. 3). Further investigation in human testes during developmental stages suggests that *LINC01016* RNA starts appearing as early as the 13^th^ year and as late as the 63^rd^ year (Fig. 3B). These results strongly suggest a potential role played by *LINC01016* plausibly in germ cell development, especially in males.

**Figure 3.**
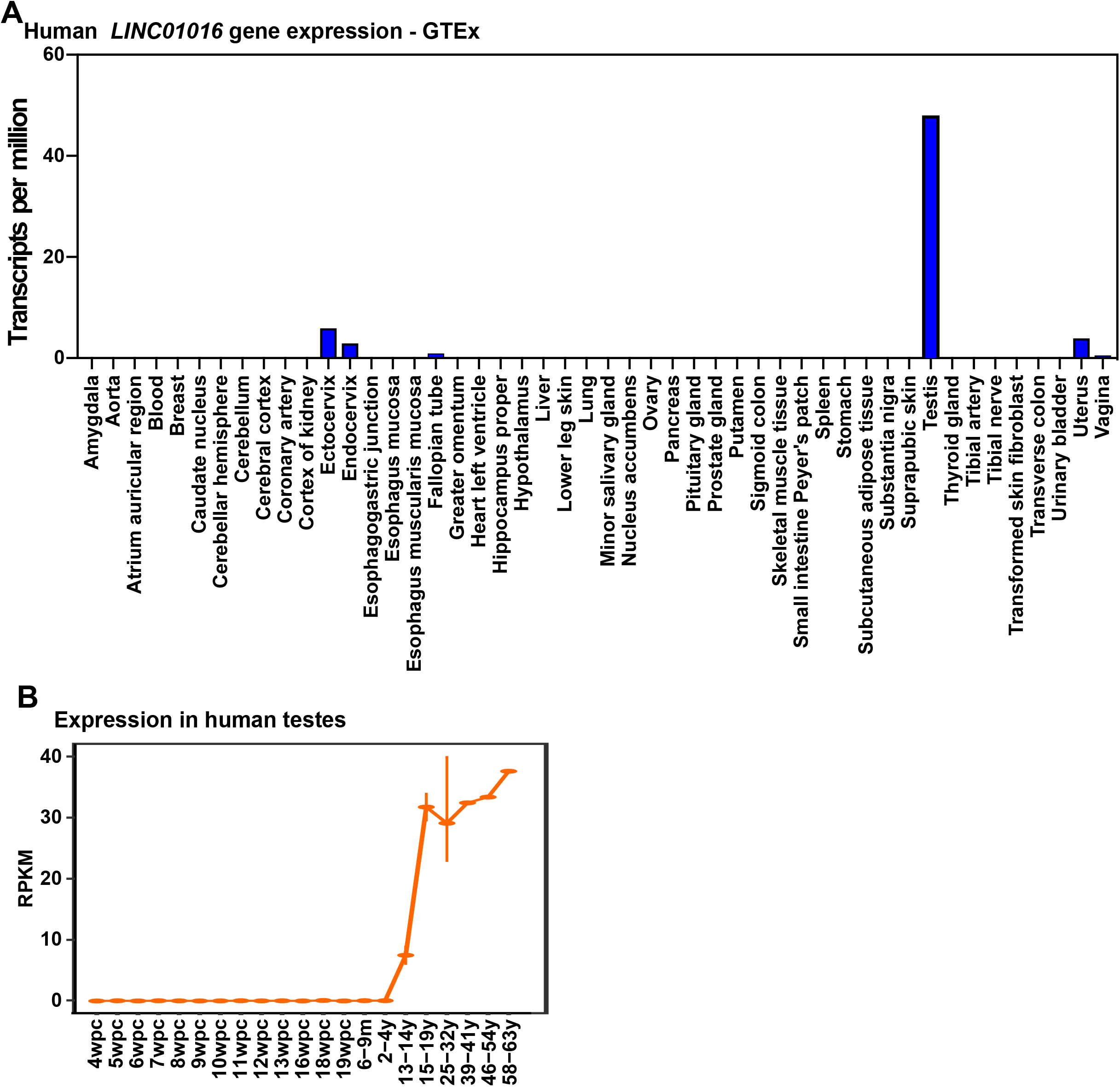
Expression of *LINC01016* in different human tissues. **A**. *LINC01016* gene expression in all the normal human tissues, as accessed from GTEx. Results are graphed as transcripts per million. **B**. *LINC01016* gene expression across developmental stages of the human testis, represented as RPKM (Reads Per Kilobase of per Million mapped reads) (wpc: weeks post-conception; m: months; y: years). The expression data were accessed using the https://apps.kaessmannlab.org/lncRNA_app/.

### *De novo* prediction of splice events, expression, and isoform identification in testis and cervix

To independently evaluate and define the *LINC01016* gene structure, RNA-Seq datasets (20-22) were obtained, and splice events were determined based on *de novo* predictions for *LINC01016* in the testis (Fig. 4A) and cervix (Fig. 5A). Testis splice graph analysis (Fig. 4A) shows 3 predicted exons and 2 junctions for eight normal testis samples. The heat map shows differential exon usage and expression (FPKM) based on each sample. On the other hand, cervix splice graph analysis (Fig. 5A) shows 4 predicted exons and 3 junctions for four normal cervix samples. The heat map shows differential exon usage and expression (FPKM) based on each sample, with the lowest expression for junction 1.

**Figure 4.**
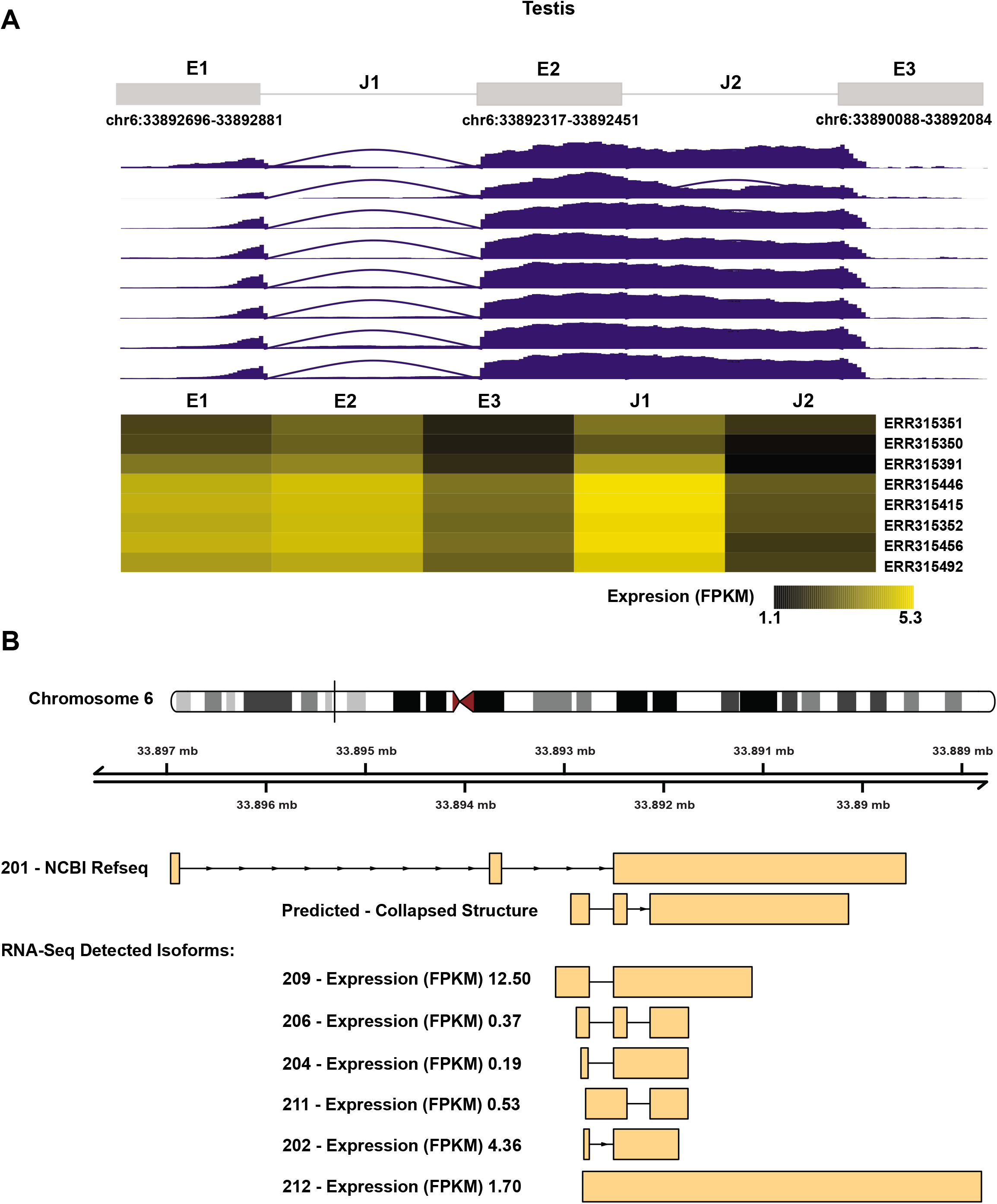
Testis RNA-Seq predicted splice events, quantification expression, and isoform identification. **A**. Splice graph analysis based on *de novo* prediction for *LINC01016* on testis samples, coverage, and expression quantification from RNA-Seq normal datasets. **B**. Gene models for *LINC01016* isoforms expression detected on RNA-Seq normal testis datasets. Included is isoform 201, which is NCBI Refseq, and the predicted collapsed structure.

**Figure 5.**
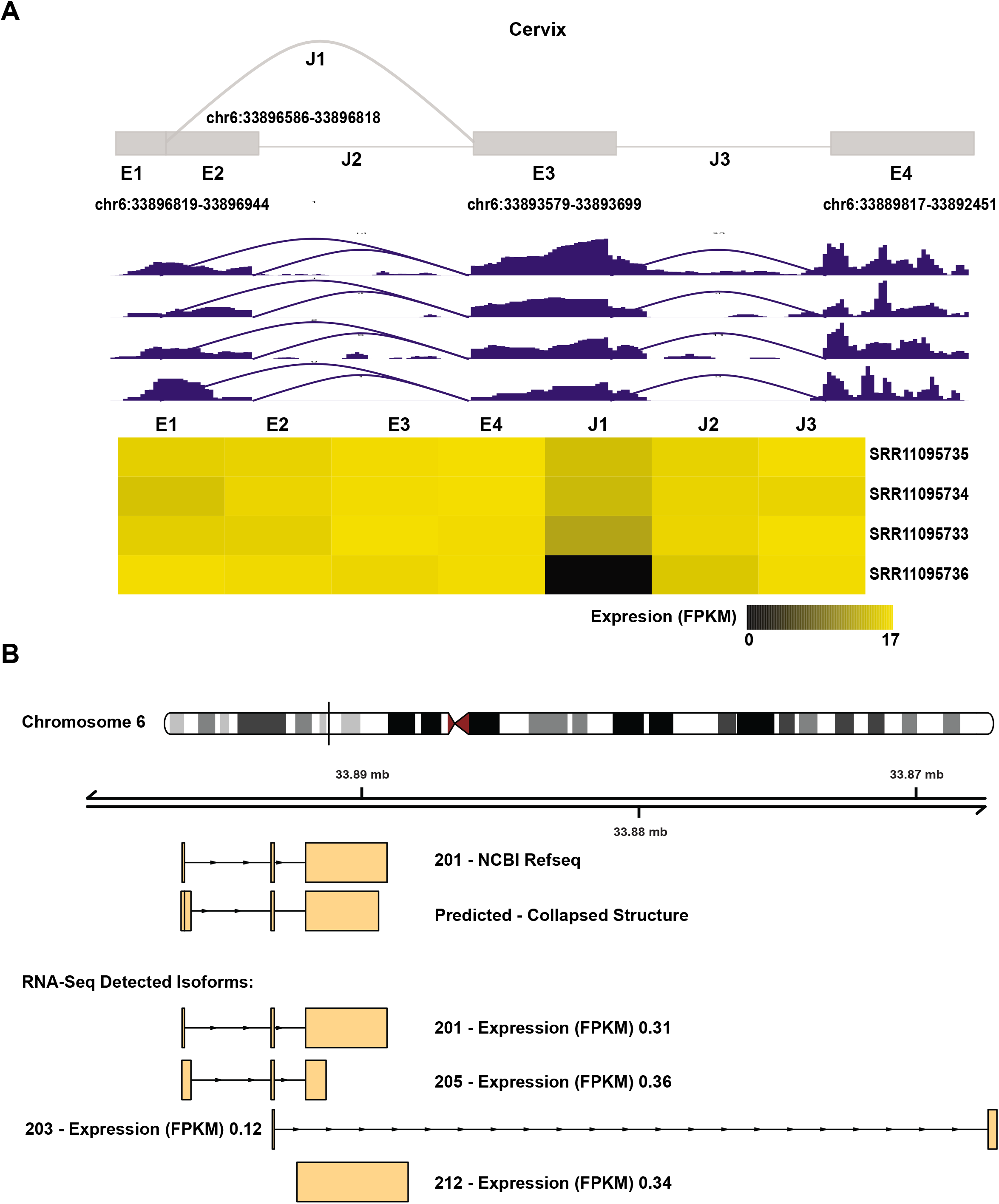
Cervix RNA-Seq predicted splice events, quantification expression, and isoform identification. **A**. Splice graph analysis based on *de novo* prediction for *LINC01016* on cervix samples, coverage, and expression quantification from RNA-Seq normal datasets. **B**. Gene models for *LINC01016* isoforms expression detected on RNA-Seq normal cervix datasets. Included is isoform 201, which is NCBI Refseq, and the predicted collapsed structure.

Gene models for the *LINC01016* isoforms detected on the RNA-Seq datasets are shown for normal testis (Fig. 4B) and cervix (Fig. 5B). The top panel shows isoform 201 (NCBI Refseq) and the predicted collapse structure for each tissue. The bottom panel depicts the different RNA-seq detected isoforms; in the case of testis, a total of 6 different isoforms were detected with an expression range of 0.19 – 12.50 FPKM (Fig. 4B), while cervix tissue analysis identified a total of 4 isoforms with an expression range of 0.12-0.36 FPKM (Fig. 5B). The gene-specific mature transcript analyses show differential exon usage and isoform expression based on tissue-specificity for normal samples.

### Comparison of *LINC01016* gene models between GTEx and RNA-Seq sample predictions

Human *LINC01016* shows diverse isoform expression depending on the tissue. However, there is variability in the expression of the isoforms of *LINC01016* depending on the data source and method used to analyze the samples. GTEx database (Release V8) shows only 2 isoforms for cervix/uterus (Fig. 6A), while our RNA-Seq *de novo* analysis predicts a total of 4 detected isoforms (Fig. 6C). Moreover, for testis, GTEx (Release V8) shows three isoforms (Fig. 6A), while our RNA-seq *de novo* analysis predicts 6 different isoforms for the samples (Fig. 6B). Importantly, our RNA-seq data analysis uses the latest release of the gene annotations for *LINC01016*, which includes a higher number of isoforms. Nevertheless, the intricate and differential expression profile of *LINC01016* transcripts indicates that each isoform is plausibly associated with distinct biological processes.

**Figure 6.**
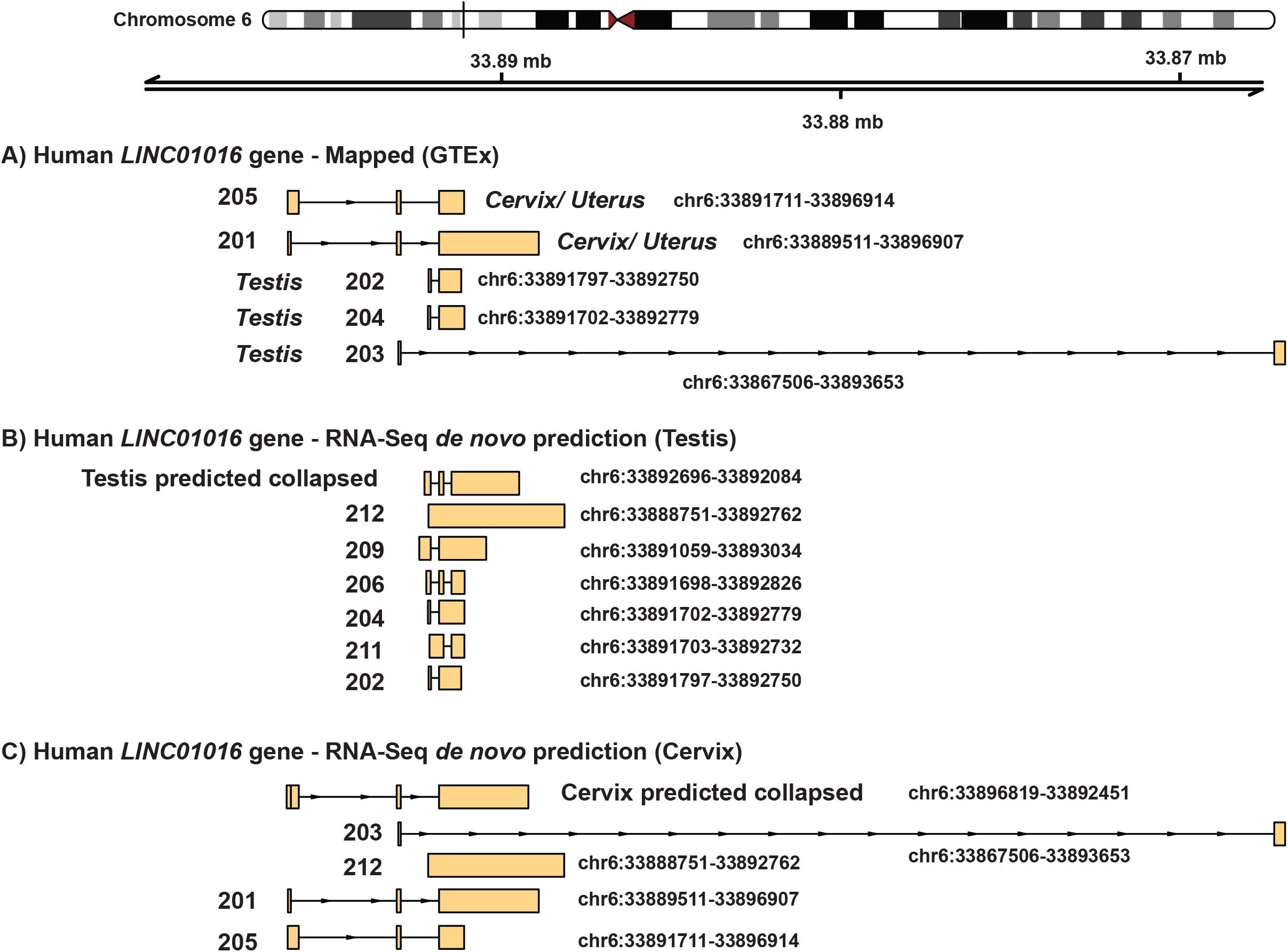
Comparison of gene models for *LINC01016* between normal GTEx, RNA-seq predicted normal testis, and RNA-seq predicted normal cervix. **A**. Isoforms detected by GTEx database v8 (dbGaP Accession phs000424.v8.p2) for normal cervix/uterus and testis samples. **B**. Gene models for *LINC01016* RNA-Seq *de novo* predicted isoforms expressed on normal testis samples. **C**. Gene models for *LINC01016* RNA-Seq *de novo* predicted isoforms expressed on normal cervix samples.

### Functional analysis of *LINC01016* isoforms

To unravel and define isoform-specific roles, six different transcripts of the *LINC01016* human gene were cloned in a mammalian expression vector and sequenced. Each transcript was expressed in a heterologous cell line (HEK293 cells, *LINC01016* is not transcribed). An advantage of this strategy is that endogenous lncRNA-driven compensatory mechanisms, which could potentially mask the isoform-specific analyses, can be avoided. Figure 7 shows the gene model diagrams of the cloned transcripts with the chromosome location, sequencing coverage, junction plots, and RNA-Seq transcript expression (FPKM) for each independent replicate. The heat map depicts the differential expression fold change for each cloned transcript (Fig. 8A). Results show significant variability in gene expression depending on the transcript. Intersections of differentially expressed genes were analyzed between samples in different groups: All regulated genes (Fig. 8B), Up-regulated genes (Fig. 8C), and Down-regulated genes (Fig. 8D). These results clearly show that although there are a common set of genes regulated by all the transcripts, the non-overlapping isoform-dependent gene sets are more extensive and mutually exclusive.

**Figure 7.**
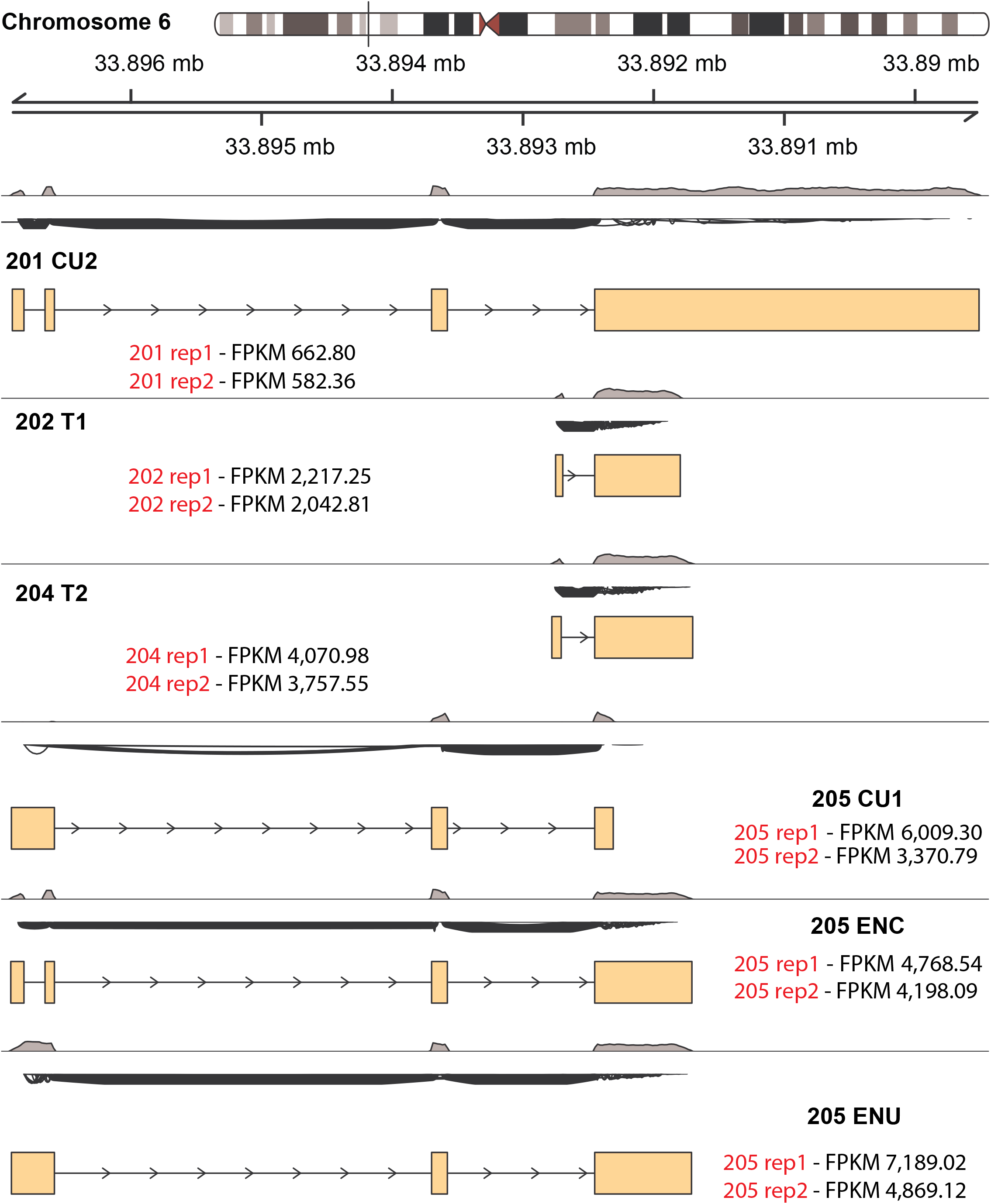
Gene models, coverage, junctions, and expression for *LINC01016* cloned transcripts on experimental RNA-Seq data. The diagram shows the chromosome location, sequencing coverage, junction plots, gene models, and RNA-Seq transcript expression (FPKM) for each cloned transcript: 201 CU2, 202 T1, 204 T2, 205 CU1, 205 ENC, and 205 ENU.

**Figure 8.**
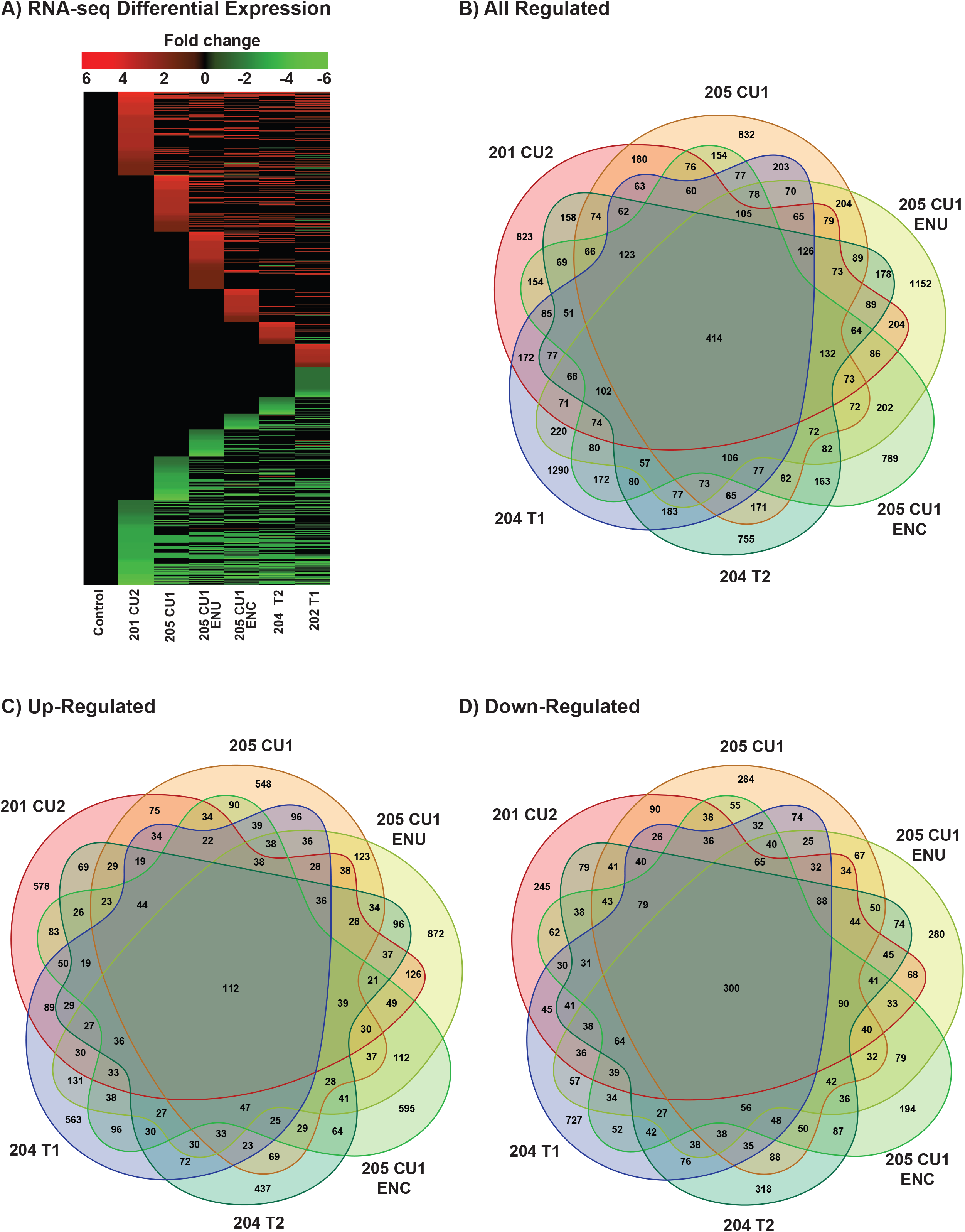
*LINC01016* cloned transcripts RNA-Seq differential expression and Venn diagrams of gene overlaps. **A**. Heatmap showing the log2(FoldChange) differential expression on cloned *LINC01016* transcripts. **B**. Venn diagrams for all regulated genes from overlapping intersections between the different categories. **C**. Venn diagrams for up-regulated genes from overlapping intersections between the different categories **D**. Venn diagrams for down-regulated genes from overlapping intersections between the different categories

### Transcript-specific molecular signatures of regulated biological pathways

To study *LINC01016* transcripts’ potential roles in biological pathways, we performed a Gene Ontology analysis. Different molecular signature databases were queried to obtain more insights into the potential transcript-specific biological functions of *LINC01016*. Heat maps showing normalized enrichment scores were generated for the different datasets: Hallmark gene sets (Fig. 9A), KEGG Pathways (Fig. 9B), molecular functions (Fig. 9C), and biological processes (Fig. 9D). All molecular signature databases showed significant variability of pathways involved depending on the selected transcript, which correlates well with the identified gene sets specific to each isoform.

**Figure 9.**
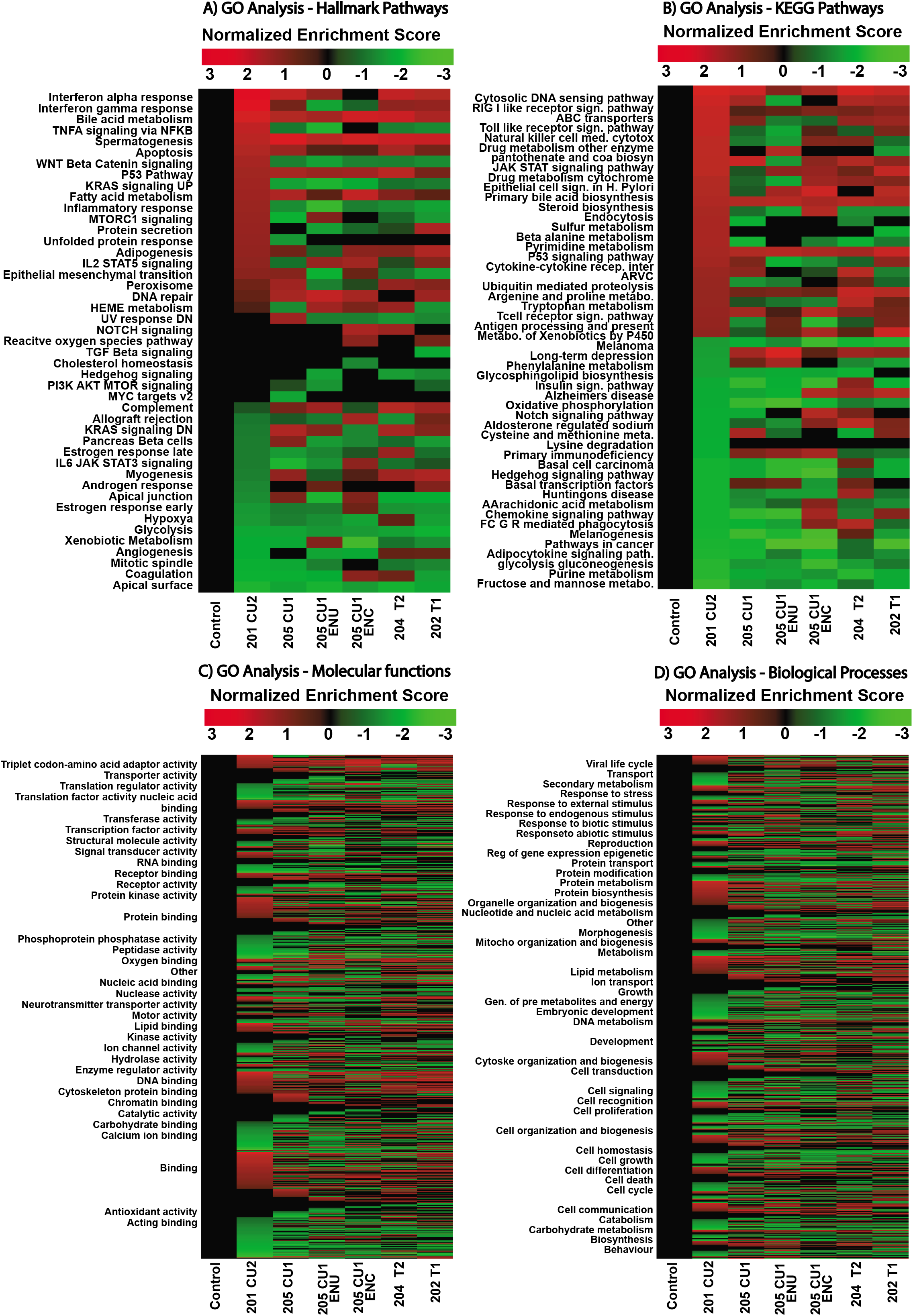
*LINC01016* cloned transcripts RNA-Seq Gene Ontology and KEGG Pathways analysis. **A**. Gene Ontology (GO) analysis for Cancer Hallmark Pathways using Normalized Enrichment Score (NES) **B**. KEGG Pathways heatmap for top 25 up/down regulated pathways on the RNA-Seq dataset using NES. **C**. GO analysis for Molecular Functions signatures using NES (molecular functions are grouped by parental term). **D**. GO analysis for Biological Processes signatures using NES (Biological Processes are grouped by parental term).

### Expression and possible prognostic value of *LINC01016* in cervical squamous cell carcinoma, testicular germ cell tumor, and endometrial carcinoma cancer

Differentially expressed lncRNAs in reproductive tissues have been thought to be potential tumor biomarkers (4), prompting us to explore the expression of *LINC01016* in malignant tissue types of cervix, testis, and uterus. By interrogating the TCGA database, we found that *LINC01016* is downregulated in cervical squamous cell carcinoma and endocervical adenocarcinoma, testicular germ cell tumor, and uterine corpus endometrial carcinoma tissues (5,23) (Fig. 10A). Further, to determine the clinical value of its altered expression patterns across these cancers, we generated Kaplan Meier plots using a cancer resource containing samples of cervical squamous cell carcinoma, testicular germ cell tumor, or uterine corpus endometrial carcinoma. We observed elevated levels of *LINC01016* RNA to be indicative of relapse-free survival in uterine corpus endometrial carcinoma and testicular germ cell tumors, but not in cervical squamous cell carcinoma (Fig. 10). Altogether, these results suggest that *LINC01016* is a testis tissue-specific lncRNA with prognostic value in a subset of cancer types.

**Figure 10.**
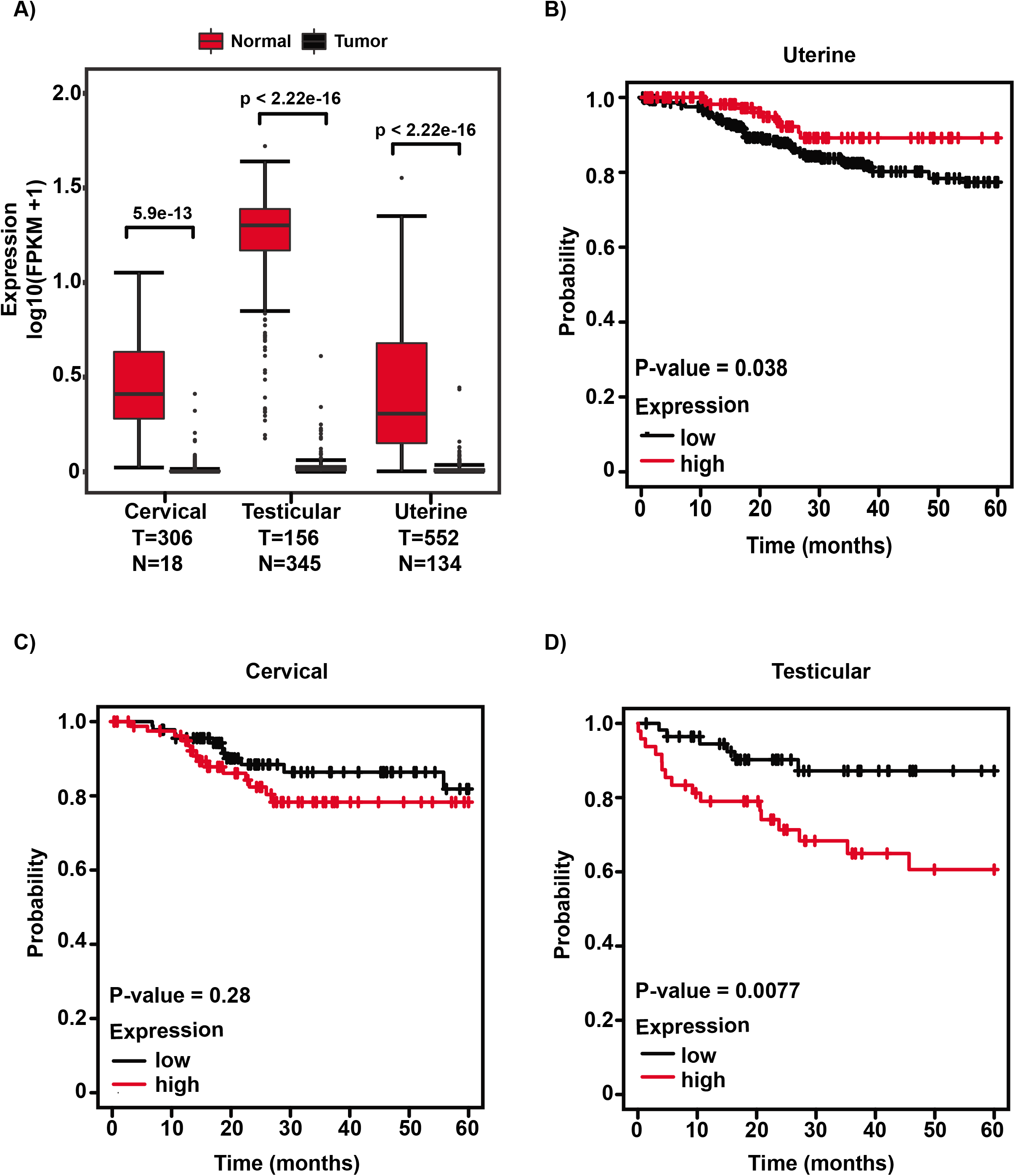
Expression of *LINC01016* in reproductive cancers. **A**. Box plot representation of *LINC01016* in testicular germ cell tumors (TGCT); cervical squamous cell carcinoma and endocervical adenocarcinoma (CESC); and uterine corpus endometrial carcinoma (UCEC). *LINC01016* RNA levels are lower in tumors (T- black box) than in normal tissues (N- red box). The expression data was accessed from the GTEx portal (Version 8) and TCGA GDC databases (v28). **B-D**. Kaplan-Meier survival analyses of patients expressing higher (red line) or lower (black line) levels of *LINC01016* RNA in uterine corpus endometrial carcinoma (**B**), cervical squamous cell carcinoma (**C**), and testicular germ cell tumors (**D**). The cancer outcome-linked gene expression data was accessed and graphed using www.kmplot.com. (19).

## Discussion

LncRNAs are found to be aberrantly expressed in tumors and can be excellent diagnostic markers with therapeutic value (1,24). Interestingly, a subset of these lncRNAs is differentially expressed spatiotemporally across reproductive tissue types such as ovary, placenta, and testis (7,24-26). While few have been investigated for their function, the lack of proper annotation presents a formidable challenge in understanding the molecular role of newly identified lncRNAs (6). Genome-wide transcriptomic analyses help define molecular features of lncRNAs, including ends of transcripts, exon usage, and relative isoform abundance (27); however, detailed gene-specific analysis remains the gold standard to accurately determine the gene structure and quantify the isoform-specific expression of relatively less abundant lncRNAs. In this regard, we have used novel and public datasets to annotate and quantify a testis-specific long intergenic noncoding RNA, *LINC01016*, in humans and non-human primates. Our results indicate that the Ensembl annotation predicts isoforms with one to four exons (Fig. 1), while similar to Ensembl data, our analysis suggests that *LINC01016* consists of four exons (Fig. 2). Contrary, exon and exon-exon expression analyses show that *LINC01016* with three exons is the predominantly expressed transcript in the cervix and uterus tissue. The signal-, space-, and time-dependent variability in the expression of isoforms is a widely observed feature of human transcripts (28). Overall, these results suggest an intricate organization of the *LINC01016* gene. This information is very intriguing since variable isoform expression through differential exon usage has been linked to several cancers (29,30). The aberrant expression of an isoform from the same gene could lead to protein or RNA with altered functions due to changes in their secondary structures or conformations (31). Hence, it is imperative to study individual isoforms of a lncRNA gene since each one of them could be functioning independently.

In addition to gene structure, *LINC01016* expression varies across tissues and organs (Fig. 3). This tissue-specific expression can be leveraged as a potential diagnostic or prognostic marker, a feature common to several lncRNAs (5,7,32). We also observed that in a manner similar to several reproductive tissue-specific genes (7), *LINC01016* is expressed during developmental stages of testis tissue, indicating a plausible role in germ cell development (33). Given the tissue specificity of *LINC01016*, it will be interesting to study its role in the testis, which could lead to potential functioning in reversibly arresting sperm production or in contraceptive technologies aimed at germ cell development. For this purpose, the majority of genes currently studied are those encoding proteins, leaving the gold mine of unstudied lncRNA genes unexplored (34), which gives an ample opportunity to explore them as novel targets.

Our data shows *LINC01016* has significant differential exon usage (Fig. 4, 5 & S1), as well as isoform-specific roles in diverse biological processes (Fig. 9). This is important because this characteristic contributes to the variability in detected isoforms across different samples and to plausible functions regulating various biological processes depending on isoform, tissue-specificity, and present conditions at a specific time point. Differential gene expression of *LINC01016* (Fig. 8) from RNA-Seq using diverse transcripts is a clear representation of gene regulation variability. Intersectional Venn diagrams from differentially expressed genes show some overlap of regulation between different isoforms of *LINC01016*, but for the most part, each transcript has a regulation of specific gene networks. Similarly, Gene Ontology analysis (Fig. 9) shows that regulation of biological pathways is isoform-dependent.

It is critical to point out the latest releases of widely used genomic databases such as GTEx, and TCGA shows only a portion of the identified *LINC01016* isoforms; 5 (Fig. 6 & S2) because their analysis is based on previous gene annotations and therefore might not reflect the full extent of known isoforms. Newer annotations from Ensembl (version 101) and GENCODE (version 37) show 12 additional isoforms of *LINC01016* matching predictions from RNA-Seq experimental sample datasets. Therefore it is crucial to compare versions and parameters of genomic databases when designing a study.

Many reproductive tissue-specific lncRNAs are differentially expressed in various cancers due to epigenetically dysregulated promoters and enhancers (2). Notably, *LINC01016* is downregulated in cervical squamous cell carcinoma, uterine corpus endometrial carcinoma samples, and in testicular germ cell tumors (Fig. 10). Survival analysis in uterine corpus endometrial carcinoma and cervical squamous cell carcinoma suggests higher *LINC01016* expression to be a predictor of a clinical outcome in patients. Previously, *LINC01016* has also been implicated in endometrial and breast cancer (35,36). Collectively, these results imply that *LINC01016* could be a promising tumor biomarker.

Our study also provides a unique perspective on using previously untapped information embedded in deep-sequencing results from species that are otherwise intractable experimental animal models (17,37). Because of the poor conservation in mice, the most widely used mammalian system, studying primate-specific lncRNA genes, is challenging due to limited comparable models in which application of lncRNA findings could be recapitulated (2). Thus, our results provide resources/experimental models as sources of information to bridge this gap in understanding, potentially aiding in developing an experimentally testable hypothesis.

In summary, through our study, we show a) complexity of gene organization at DNA and transcript level can be understood by integrating publicly available data with targeted experimental approaches, b) differential exon usage, c) evolutionary significance of *LINC01016*, d) relative expression of *LINC01016* in tissues with a plausible role in reproductive physiology, e) isoform-dependent differences in regulation of gene networks and biological pathways, and f) *LINC01016* as a potential tumor biomarker.

## Acknowledgments

We thank Alana L. Harrison and members of the Gadad lab for their helpful comments. We thank Texas Tech University High-Performance Computing Center (Lubbock) and TTUHSC El Paso NGS Computing Server (El Paso). S.S.G. is a CPRIT Scholar in Cancer Research. This work was supported by a first-time faculty recruitment award from the Cancer Prevention and Research Institute of Texas (CPRIT; RR170020).

## Author contributions

Enrique Ramos – data curation, methodology, formal analysis; Barbara Yang – data curation; Yasmin M. Vasquez – investigation, writing - review & editing; Ramesh Choudhari – data curation; Ken Y. Lin – writing, investigation, review & editing; Shrikanth S. Gadad – conceptualization, methodology, investigation, funding acquisition, project administration, investigation, writing - original draft, writing - review & editing

## Figure Legends

**Figure S1.**
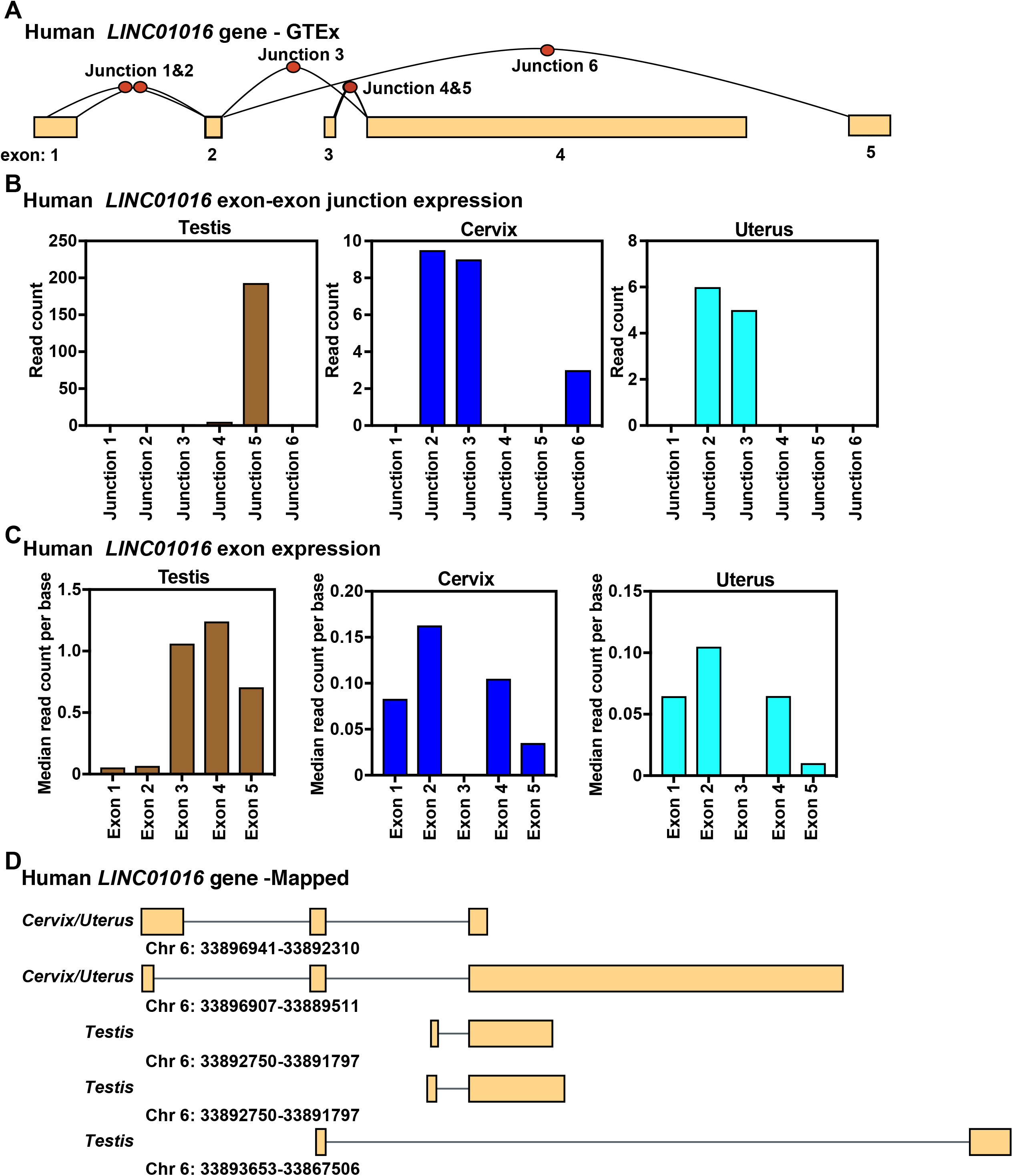
Characterization of the structure of the human *LINC01016* gene by analysis of RNA-Seq data. A. Collapsed view of human *LINC01016* gene exon and exon-exon junction structure from GTEx (8,9). **B**. Exon-exon junction levels in testes expressed as read count, accessed from GTEx. **C**. Exon-specific expression in testes obtained from GTEx testes. **D**. Diagram of the human *LINC01016* isoforms as analyzed in parts A-C above. Exons are depicted as boxes and introns as lines.

**Figure S2.**
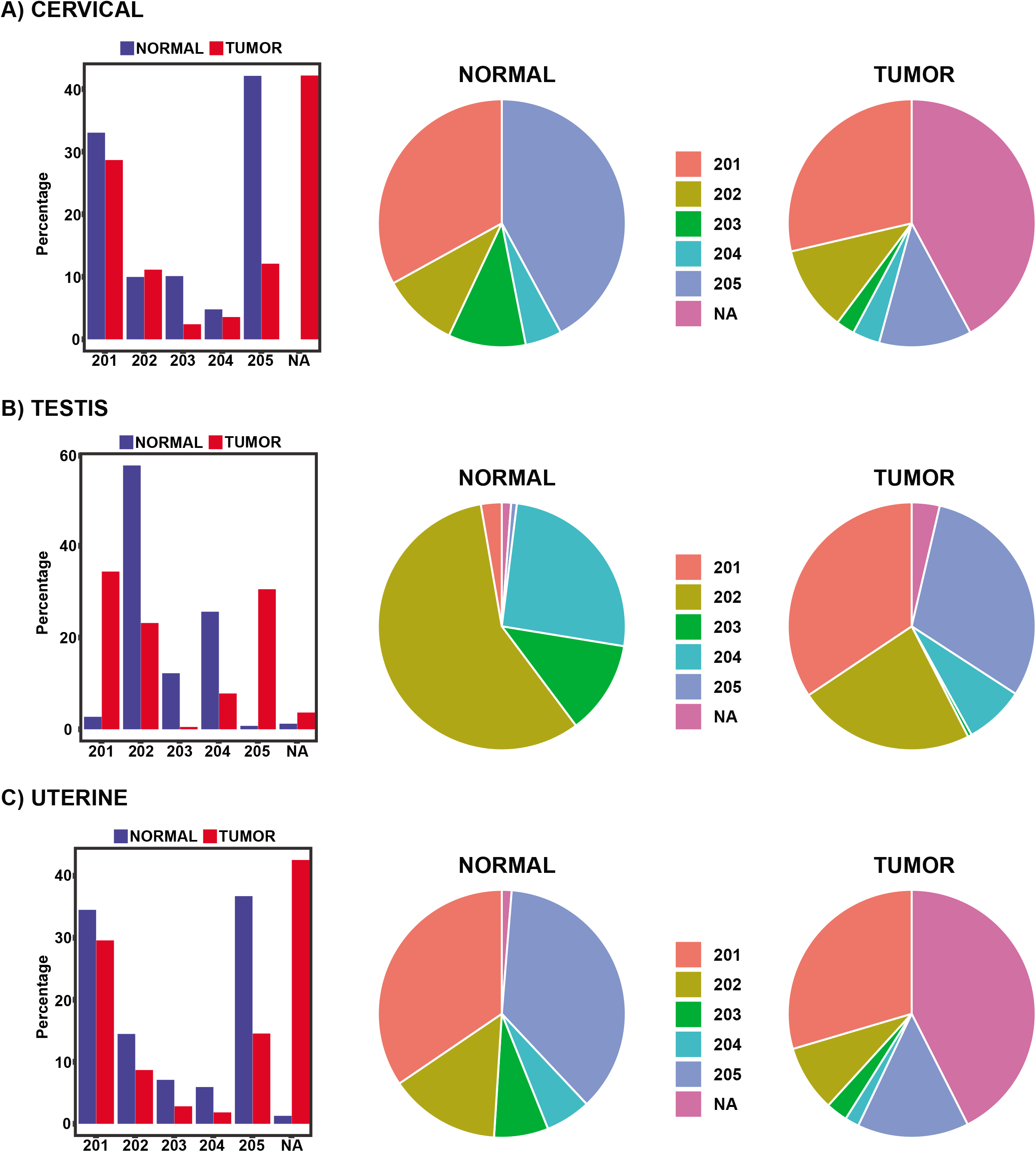
GTEx and TCGA *LINC01016* transcript expression from available RNA-Seq datasets on different tissues. *LINC01016* isoforms detected by GTEx database (8,9) for normal tissue and TCGA Pan-Cancer (PANCAN) tumor samples for **A**. Cervical, **B**. Testis, and **C**. Uterine tissue samples. Isoform annotation is based on GENCODE v22.

## References

1. Camacho CV, Choudhari R, Gadad SS. Long noncoding RNAs and cancer, an overview. Steroids. 2018;133:93–95.

2. Choudhari R, Sedano MJ, Harrison AL, Subramani R, Lin KY, Ramos EI, Lakshmanaswamy R, Gadad SS. Long noncoding RNAs in cancer: From discovery to therapeutic targets. Adv Clin Chem. 2020;95:105–147.

3. Sun M, Gadad SS, Kim DS, Kraus WL. Discovery, Annotation, and Functional Analysis of Long Noncoding RNAs Controlling Cell-Cycle Gene Expression and Proliferation in Breast Cancer Cells. Mol Cell. 2015;59(4):698–711.

4. Hosono Y, Niknafs YS, Prensner JR, Iyer MK, Dhanasekaran SM, Mehra R, Pitchiaya S, Tien J, Escara-Wilke J, Poliakov A, Chu SC, Saleh S, Sankar K, Su F, Guo S, Qiao Y, Freier SM, Bui HH, Cao X, Malik R, Johnson TM, Beer DG, Feng FY, Zhou W, Chinnaiyan AM. Oncogenic Role of THOR, a Conserved Cancer/Testis Long Non-coding RNA. Cell. 2017;171(7):1559–1572 e1520.

5. Vasquez YM, Nandu TS, Kelleher AM, Ramos EI, Gadad SS, Kraus WL. Genome-wide analysis and functional prediction of the estrogen-regulated transcriptional response in the mouse uterusdagger. Biol Reprod. 2020;102(2):327–338.

6. Uszczynska-Ratajczak B, Lagarde J, Frankish A, Guigo R, Johnson R. Towards a complete map of the human long non-coding RNA transcriptome. Nat Rev Genet. 2018;19(9):535–548.

7. Choudhari R, Yang B, Rotwein P, Gadad SS. Structure and expression of the long noncoding RNA gene MIR503 in humans and non-human primates. Mol Cell Endocrinol. 2020:110819.

8. e GP. Enhancing GTEx by bridging the gaps between genotype, gene expression, and disease. Nat Genet. 2017;49(12):1664–1670.

9. Consortium GT, Laboratory DA, Coordinating Center - Analysis Working G, Statistical Methods groups-Analysis Working G, Enhancing Gg, Fund NIHC, Nih/Nci, Nih/Nhgri, Nih/Nimh, Nih/Nida, Biospecimen Collection Source Site N, Biospecimen Collection Source Site R, Biospecimen Core Resource V, Brain Bank Repository-University of Miami Brain Endowment B, Leidos Biomedical-Project M, Study E, Genome Browser Data I, Visualization EBI, Genome Browser Data I, Visualization-Ucsc Genomics Institute UoCSC, Lead a, Laboratory DA, Coordinating C, management NIHp, Biospecimen c, Pathology, e QTLmwg, Battle A, Brown CD, Engelhardt BE, Montgomery SB. Genetic effects on gene expression across human tissues. Nature. 2017;550(7675):204–213.

10. Quintana-Murci L. Understanding rare and common diseases in the context of human evolution. Genome Biol. 2016;17(1):225.

11. Manolio TA, Fowler DM, Starita LM, Haendel MA, MacArthur DG, Biesecker LG, Worthey E, Chisholm RL, Green ED, Jacob HJ, McLeod HL, Roden D, Rodriguez LL, Williams MS, Cooper GM, Cox NJ, Herman GE, Kingsmore S, Lo C, Lutz C, MacRae CA, Nussbaum RL, Ordovas JM, Ramos EM, Robinson PN, Rubinstein WS, Seidman C, Stranger BE, Wang H, Westerfield M, Bult C. Bedside Back to Bench: Building Bridges between Basic and Clinical Genomic Research. Cell. 2017;169(1):6–12.

12. Soumillon M, Cacchiarelli D, Semrau S, van Oudenaarden A, Mikkelsen TS. Characterization of directed differentiation by high-throughput single-cell RNA-Seq. bioRxiv. 2014:003236.

13. Vera M, Biswas J, Senecal A, Singer RH, Park HY. Single-Cell and Single-Molecule Analysis of Gene Expression Regulation. Annu Rev Genet. 2016;50:267–291.

14. Sarropoulos I, Marin R, Cardoso-Moreira M, Kaessmann H. Developmental dynamics of lncRNAs across mammalian organs and species. Nature. 2019;571(7766):510–514.

15. Peng X, Thierry-Mieg J, Thierry-Mieg D, Nishida A, Pipes L, Bozinoski M, Thomas MJ, Kelly S, Weiss JM, Raveendran M, Muzny D, Gibbs RA, Rogers J, Schroth GP, Katze MG, Mason CE. Tissue-specific transcriptome sequencing analysis expands the non-human primate reference transcriptome resource (NHPRTR). Nucleic Acids Res. 2015;43(Database issue):D737–742.

16. Choudhari R, Yang B, Rotwein P, Gadad SS. Structure and expression of the long noncoding RNA gene MIR503 in humans and non-human primates. Mol Cell Endocrinol. 2020;510:110819.

17. Rotwein P. The insulin-like growth factor 2 gene and locus in nonmammalian vertebrates: Organizational simplicity with duplication but limited divergence in fish. J Biol Chem. 2018;293(41):15912–15932.

18. Tang Z, Li C, Kang B, Gao G, Li C, Zhang Z. GEPIA: a web server for cancer and normal gene expression profiling and interactive analyses. Nucleic Acids Res. 2017;45(W1):W98–W102.

19. Nagy A, Lanczky A, Menyhart O, Gyorffy B. Validation of miRNA prognostic power in hepatocellular carcinoma using expression data of independent datasets. Sci Rep. 2018;8(1):9227.

20. Djureinovic D, Fagerberg L, Hallstrom B, Danielsson A, Lindskog C, Uhlen M, Ponten F. The human testis-specific proteome defined by transcriptomics and antibody-based profiling. Mol Hum Reprod. 2014;20(6):476–488.

21. Xu J, Zou J, Wu L, Lu W. Transcriptome analysis uncovers the diagnostic value of miR-192-5p/HNF1A-AS1/VIL1 panel in cervical adenocarcinoma. Sci Rep. 2020;10(1):16584.

22. Xu J, Zhang Y, Huang Y, Dong X, Xiang Z, Zou J, Wu L, Lu W. circEYA1 Functions as a Sponge of miR-582-3p to Suppress Cervical Adenocarcinoma Tumorigenesis via Upregulating CXCL14. Mol Ther Nucleic Acids. 2020;22:1176–1190.

23. Cancer Genome Atlas Research N, Weinstein JN, Collisson EA, Mills GB, Shaw KR, Ozenberger BA, Ellrott K, Shmulevich I, Sander C, Stuart JM. The Cancer Genome Atlas Pan-Cancer analysis project. Nat Genet. 2013;45(10):1113–1120.

24. Vallone C, Rigon G, Gulia C, Baffa A, Votino R, Morosetti G, Zaami S, Briganti V, Catania F, Gaffi M, Nucciotti R, Costantini FM, Piergentili R, Putignani L, Signore F. Non-Coding RNAs and Endometrial Cancer. Genes (Basel). 2018;9(4).

25. Taylor DH, Chu ET, Spektor R, Soloway PD. Long non-coding RNA regulation of reproduction and development. Mol Reprod Dev. 2015;82(12):932–956.

26. Wang Q, Wang N, Cai R, Zhao F, Xiong Y, Li X, Wang A, Lin P, Jin Y. Genome-wide analysis and functional prediction of long non-coding RNAs in mouse uterus during the implantation window. Oncotarget. 2017;8(48):84360–84372.

27. Trapnell C, Williams BA, Pertea G, Mortazavi A, Kwan G, van Baren MJ, Salzberg SL, Wold BJ, Pachter L. Transcript assembly and quantification by RNA-Seq reveals unannotated transcripts and isoform switching during cell differentiation. Nat Biotechnol. 2010;28(5):511–515.

28. Reyes A, Huber W. Alternative start and termination sites of transcription drive most transcript isoform differences across human tissues. Nucleic Acids Res. 2018;46(2):582–592.

29. Robinson JT, Thorvaldsdottir H, Wenger AM, Zehir A, Mesirov JP. Variant Review with the Integrative Genomics Viewer. Cancer Res. 2017;77(21):e31–e34.

30. Zhang Z, Pal S, Bi Y, Tchou J, Davuluri RV. Isoform level expression profiles provide better cancer signatures than gene level expression profiles. Genome Med. 2013;5(4):33.

31. Tian N, Li J, Shi J, Sui G. From General Aberrant Alternative Splicing in Cancers and Its Therapeutic Application to the Discovery of an Oncogenic DMTF1 Isoform. Int J Mol Sci. 2017;18(3).

32. Cabili MN, Trapnell C, Goff L, Koziol M, Tazon-Vega B, Regev A, Rinn JL. Integrative annotation of human large intergenic noncoding RNAs reveals global properties and specific subclasses. Genes Dev. 2011;25(18):1915–1927.

33. Deng X, Berletch JB, Nguyen DK, Disteche CM. X chromosome regulation: diverse patterns in development, tissues and disease. Nat Rev Genet. 2014;15(6):367–378.

34. Kent K, Johnston M, Strump N, Garcia TX. Toward Development of the Male Pill: A Decade of Potential Non-hormonal Contraceptive Targets. Front Cell Dev Biol. 2020;8:61.

35. Jonsson P, Coarfa C, Mesmar F, Raz T, Rajapakshe K, Thompson JF, Gunaratne PH, Williams C. Single-Molecule Sequencing Reveals Estrogen-Regulated Clinically Relevant lncRNAs in Breast Cancer. Mol Endocrinol. 2015;29(11):1634–1645.

36. Pan X, Li D, Huo J, Kong F, Yang H, Ma X. LINC01016 promotes the malignant phenotype of endometrial cancer cells by regulating the miR-302a-3p/miR-3130-3p/NFYA/SATB1 axis. Cell Death Dis. 2018;9(3):303.

37. Rotwein P. Quantifying promoter-specific Insulin-like Growth Factor 1 gene expression by interrogating public databases. Physiol Rep. 2019;7(1):e13970.

38. Goldman MJ, Craft B, Hastie M, Repecka K, McDade F, Kamath A, Banerjee A, Luo Y, Rogers D, Brooks AN, Zhu J, Haussler D. Visualizing and interpreting cancer genomics data via the Xena platform. Nat Biotechnol. 2020;38(6):675–678.

